# Dynamics of Soil Nematode Communities Revealed Significant Variation in Greenhouse with Different Continuous Cropping Years

**DOI:** 10.1101/593541

**Authors:** Xueliang Tian, Xiaoman Zhao, Zhenchuan Mao, Bingyan Xie

## Abstract

Continuous cropping usually resulted in a series of soil problems, especially root knot nematode disaster in greenhouse because of imbalance in soil nematode community. However, the dynamic change in soil nematode community in greenhouse with various continuous cropping years during plant growing season is unclear. Here, soil nematode communities in greenhouse after 2 and 10 years of continuous cropping were comparatively analyzed using 18S rDNA high-throughput sequencing to reveal the changing processes in soil nematode communities. Compared to the 2-year greenhouse, soil in the 10-year greenhouse showed soil acidification, nutrients accumulation, salinization. In term of α-diversity, Bacterial-feeding nematodes (BF) were dominant in the 2-year greenhouse over the whole growing season, but plant-parasitic nematodes (PP) became the dominant group in the 10-year greenhouse in the late growing season. *Meloidogyne* gradually became the dominant group (Relative abundance 70.9%) in the 10-year greenhouse, while *Rhabditidae* (Relative abundance from 99.8% to 26.8%) was the predominant members in the 2-year greenhouse. For β-diversity, hierarchical clustering analysis, unWeighted UniFrac principal component analysis (PCA) and principal co-ordinates analysis (PCoA) all revealed that soil nematode communities in the two types of greenhouses exhibited significant differences. Redundancy analysis (RDA) showed that soil nematode communities in the 10-year greenhouse were related to high soil organic material, total nitrogen, electrical conductivity and disease index of root knot nematode. Fisher’ exact test and Pearson correlation coefficients revealed that *Meloidogyne* caused main differences in soil nematode communities between the two types of greenhouses. Meanwhile, population dynamics of *Meloidogyne* were divided into dormant phase, low-level increasing phase and exponential phase during the whole season. In total, the significant variation led different dynamics of soil nematode communities in the 2-year and 10-year greenhouse.

## Introduction

Since the1980s, greenhouses have been applied in northern China because of the low costs of construction and operation, making them very economical. As an extensive and unique crop production system, they have become the main agricultural facilities in China. Currently, China has the most agricultural greenhouses in the world. Vegetables cultivated in greenhouses are the primary plantings in China [1], and they occupy 3.44 million hectares in China, which accounted for more than 90% of the world total in 2010 [2]. The solar greenhouse system provides a great deal of vegetables annually in China and increases farming incomes. However, this planting system, with its continuous cropping and short rotations, has drawbacks [3], including soil nutrient imbalances [4], soil secondary salination [5], secondary metabolite autotoxicity [6], rhizosphere microorganism disorders [7] and replanting-associated diseases, especially soil-borne disease such as root rot and wilt, as well as root-knot nematodes [8]. Finally, these drawbacks disrupt the balance of the soil ecosystem and cause retarded crop growth and nutritional disturbances, and yield and quality decline, resulting in a restriction in the development of greenhouse vegetables in China [9,10].

As the most abundant group of soil fauna, soil nematodes play important roles in the soil ecosystem, including the decay of organic materials and nutrient transformation. Because soil nematodes are sensitive to agricultural management activities [11-13], nutrient states and the soil environment [14], their populations and community structure are affected by soil disturbances. Therefore, soil nematodes are suitable indicators of soil health. Although the soil nematode communities from farmlands, forests and grasslands have been comprehensively investigated [15-17], soil nematode communities in greenhouses have been mainly identified using morphology and/or feeding habit-based classifications [1,18,19].To overcome drawback of the low efficiency of traditional morphology-based analysis, high-throughput sequencing has been used as a powerful tool for the taxonomic identification of eukaryotes and/or for their community analyses in various ecosystems. Recently, some studies have analyzed the soil nematode community using high-throughput sequencing to reveal the structure and composition of soil nematode community in various ecological soil [20-22].

Here, we assessed the soil nematode community in greenhouses with various continuous cropping years by high-throughput sequencing. The objectives of the study were (i) to elucidate differences in the dynamic of soil nematode communities during the whole growing season in greenhouse with various continuous cropping years and (ii) to reveal population dynamic of root knot nematode in greenhouse with different disease severity. For this purpose, greenhouse with 2 and 10 years cultivation period were sampled for 7 times within the whole growing season according to local planting habit and growth rhythm of cucumber.

## Materials and methods

### Site descriptions

The experiment was conducted in six typical greenhouses (three greenhouses with 2 years planting time and three greenhouses with 10 years planting time) located in Xinxiang, Henan, China (35.2’N, 113.8’E). The average annual temperature is ∼14 °C, and the average precipitation is 656.3mm. The soil type of the greenhouses is cinnamon soil according to China’s soil classification retrieval system.

The soil pH value was measured by acidometer. Soil organic matter content was assayed by burning dried soil in a muffle furnace at 490 °C for 8 h. Total N was measured colorimetrically following micro-Kjeldahl digestion. Soil organic matter was measured using the oxidation of potassium dichromate with a 0.4 mol L-1 K_2_Cr_2_O_7_-H_2_SO_4_ solution. Electric conductivity was determined by a conductivity meter in a 1:5 soil: water solution (w/v).

The six greenhouses were located within 0.5 kilometers and used for planting cucumber in the first half of the year and tomato in the latter half of the year. The 2-year greenhouse was planted for 2 years after this field was planted for wheat and maize. The 10-year greenhouse was planted for 10 years with cucumber and tomato. Each greenhouse area was ∼1,000 mm^2^, holding 50 plant beds. Cucumber seedlings were planted in two rows per bed with 20 cm between plants. Three plots were randomly set in one greenhouse. Each plot was about 300 mm^2^, contained 15 plant beds.

### Sampling

To monitor the dynamics of soil nematode populations, soil samples were taken seven times from the greenhouses in 2015: Feb. 16th (Pre-planted period), Mar. 14th (Seedling period), Apr. 3rd (Fruiting early period), Apr. 19th (Fruit harvest period), May.7th (Fruit harvest period), May.26th (Fruit harvest period) and Jun.14th (Post-harvest). Meanwhile, properties of soil sample were measured using method aforementioned.

A total of three sub-plots containing 15 beds with approximately 450 cucumber plants were arranged in each greenhouse. Each soil sample consisted of 10 soil cores (25-mm diameter× 20-cm depth) that were collected from each sub-plot with a soil auger (diameter, 25 mm) using the five-point sampling method. The sample sites were selected with approximately 5 cm from the nearest cucumber plant to a depth of 20 cm. Then, 5 cm of surface soil was removed. Samples from the same sub-plots were pooled together for each sampling time, mixed and packed into individual plastic bags. The samples were immediately taken to the laboratory.

The disease index of root knot nematode on cucumber was investigated since cucumber planting. Cucumber roots were carefully removed from the soil, and the residual soil on roots was shaken off. A root gall index (1-5) was calculated as follows: 1, no galls; 2, 1%-25% of roots with galls; 3, 26%-50% with galls; 4, 51%-75% with galls; and 5, >75% with galls [23].

### Soil nematode isolation and DNA extraction

Soil nematodes were extracted from 100 g of fresh soil per sample using the Baermann funnel method with modifications [24]. Soil nematodes were isolated for three times from each soil sample and mixed together. The soil nematodes were collected into 2.0-ml centrifuge tubes for DNA extraction. To avoid the effects of humic acid on nematode DNA, the DNA samples were extracted using the MOBIO UltraClean™ Soil DNA Isolation Kit (Mo Bio Laboratories, Carlsbad, CA, USA) according to the manufacturer’s instructions. Soil nematode DNA from 3 replicates were pooled, resulting in 42 DNA samples (7 samples taken 7 times from both three 2-years and three 10-year greenhouse).

### PCR amplification and pyrosequencing

For cost reduction, DNA sampes from same types greenhouse at same sampling time were pooled together. To generate amplicons for MiSeq pyro-sequencing, primers NF1(5′-GGT GGT GCA TGG CCG TTC TTA GTT-3′) and18Sr2b (5′-AGC GAC GGG CGG TGT GTA CAA A-3′) were used in this study. The forward primer NF1was used previously by Porazinska et al [25]. The reverse primer 18Sr2b is derived from a primer used by Darby et al [20].The PCR procedure was as follows: denaturation step of 95°C for 3 min, 35 cycles at 95°C for 30 s, 55°C for 30 s and 72°C for 45s, followed by a final elongation at 72°C for 10 min. The final volume of the PCR reactions was 20 μl, including 4 μl 5×FastPfu buffer, 2 μl 2.5 mM dNTPs, 0.8 μl forward primer (5 μM), 0.8 μl reverse primer (5 μM), 0.4 μl FastPfu polymerase, 1μl template DNA (about 10 ng) and 10.8 μl dd H_2_O. The concentration of PCR products was quantified by QuantiFluor-ST (Promega Corporation, WI, USA) according to the manufacturer’s instructions, and were then proportionally mixed according to each sample’s concentration. The sample pool was sequenced using Illumina Miseq PE300 (Illumina) by the Majorbio Company (Shanghai, China).

### Data analysis

Raw sequences were processed using QIIME software (1.7.0) [26]. Amplicon Noise and Perseus were used to remove reads mismatching the primer and barcode sequences, PCR-based or sequencing errors and chimeras [27]. Uclust was used for screening operational taxonomic units (OTUs) at 97% similarity [28]. The sequences of each OTU were aligned to the reference set of nematode sequences [21], non-nematode sequences were removed. Rarefaction curves with average number of observed OTUs were generated to compare relative levels of OTU diversity across the two greenhouse soils.

For α-diversity, the Shannon index and Simpson index was calculated to assess the diversity soil nematode community according to formula 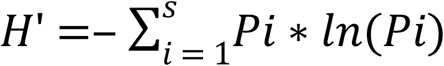 and 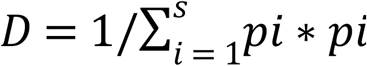, where pi is the proportion of each taxon (or OTU) in the soil nematode community, s is the total number of OTU in the soil nematode community. Based on their OTU’s taxonomy, soil nematodes were classified into bacterial feeders, fungal feeders, omnivores/predators and plant parasites [29]. Maturity index (MI) for non-plant parasites and plant-parasite index (PPI) for plant parasites was evaluated environmental disturbance of greenhouse soil, 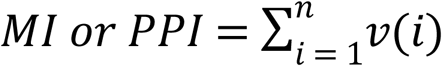, where v(i) is the c-pvalue of taxon i, and f(i) is the frequency of that taxon in a community [30]. Analysis of variance (ANOVA) was performed on Shannon index, Simpson index, MI and PPI across the two types of greenhouses at the same sampling time.

For β-diversity, hierarchical cluster dendrograms (with BrayCurtis distance dissimilarities) were performed using Mothur based on the OTU composition for comparing the soil nematode community structures across all soil samples[31]. An unWeighted UniFrac principal component analysis (PCA) and unWeighted UniFrac principal co-ordinates analysis (PCoA) were performed with Canoco to compare soil nematode communities in the 2-year and 10-year greenhouse [32]. In order to reveal the relationships between soil nematode community compositions and environmental variables, redundancy analysis (RDA) was performed according to Chen’ s method [33]. Pearson correlation bubble diagram of soil nematode trophic groups, the most ten OTUs and soil properties, root knot nematode disease index were all calculated by IBM SPSS Statistics 19.0 (SPSS, Inc., Chicago, IL, United States).

## Results

### Soil properties in greenhouses with different planting ages

As shown in S1 Table, pH in the 10-year greenhouse was markedly lower than the 2-year greenhouse (F=31.17, *P* <0.01), reflecting that soil in the 10-year greenhouse showed acidified trend. SOM and TN in the 10-year greenhouse were significantly greater than that in the 2-year greenhouse (F=8.76, *P* =0.012; F=5.55, *P* =0.036), indicating that nutrients accumulation in the 10-year greenhouse soil occurred. Likely, EC in the 10-year greenhouse were clearly higher than that in the 2-year greenhouse (F=5.13, *P* =0.043), suggesting that salinization appeared in the 10-year greenhouse. Overall, there were significant differences in soil properties between the 2-year and 10-year greenhouses.

#### Basic information of sequences

At least 10,000 raw sequences were obtained from each sample using the Illumina Miseq PE3000 platform, and the lengths of the reads were 354–360 bp (S2 Table). Sequence data have been deposited in the NCBI Sequence Read Archive (SRA) database with the accession number SRP148523. Although a soil nematode enrichment step in this study was applied to decrease biases, other groups of organisms, such as fungi, algae and plants, were amplified, and these non-nematode sequences were removed. The remaining nematode sequences were analyzed to reveal the composition of soil nematode community. The number of soil nematode sequences varied among samples, ranging from 10, 455 to 20, 388.

### Soil nematode community α-diversity

Rarefaction curves analysis of 14 samples in the two types greenhouses confirmed that the number of observed OTUs increased asymptotically with an increase in reads (Fig. 1A). In addition, the number of observed OTUs in early samples (2-1 and 10-1) less than in later samples (10-7 and 2-7). Shannon index and Simpson index showed significant differences between the two types of greenhouses on some sampling times (Fig. 1B and 1C). Shannon index in the 2-year greenhouse were markedly higher (3-14, 4-3, 4-19, 5-26 and 6-14) but significantly lower (2-16 and 5-7) than in the 10-year greenhouse (Fig 1B). Likely, Simpson index in the 2-year greenhouse were markedly higher (3-14, 4-3, 4-19 and 6-14), but significantly lower (2-16 and 5-7) than in the 10-year greenhouse (Fig 1C). Additionally, Shannon index and Simpson index in one greenhouse shifted significantly among seven sampling times (paired *t* test, *P* < 0.05).

**Fig 1.**
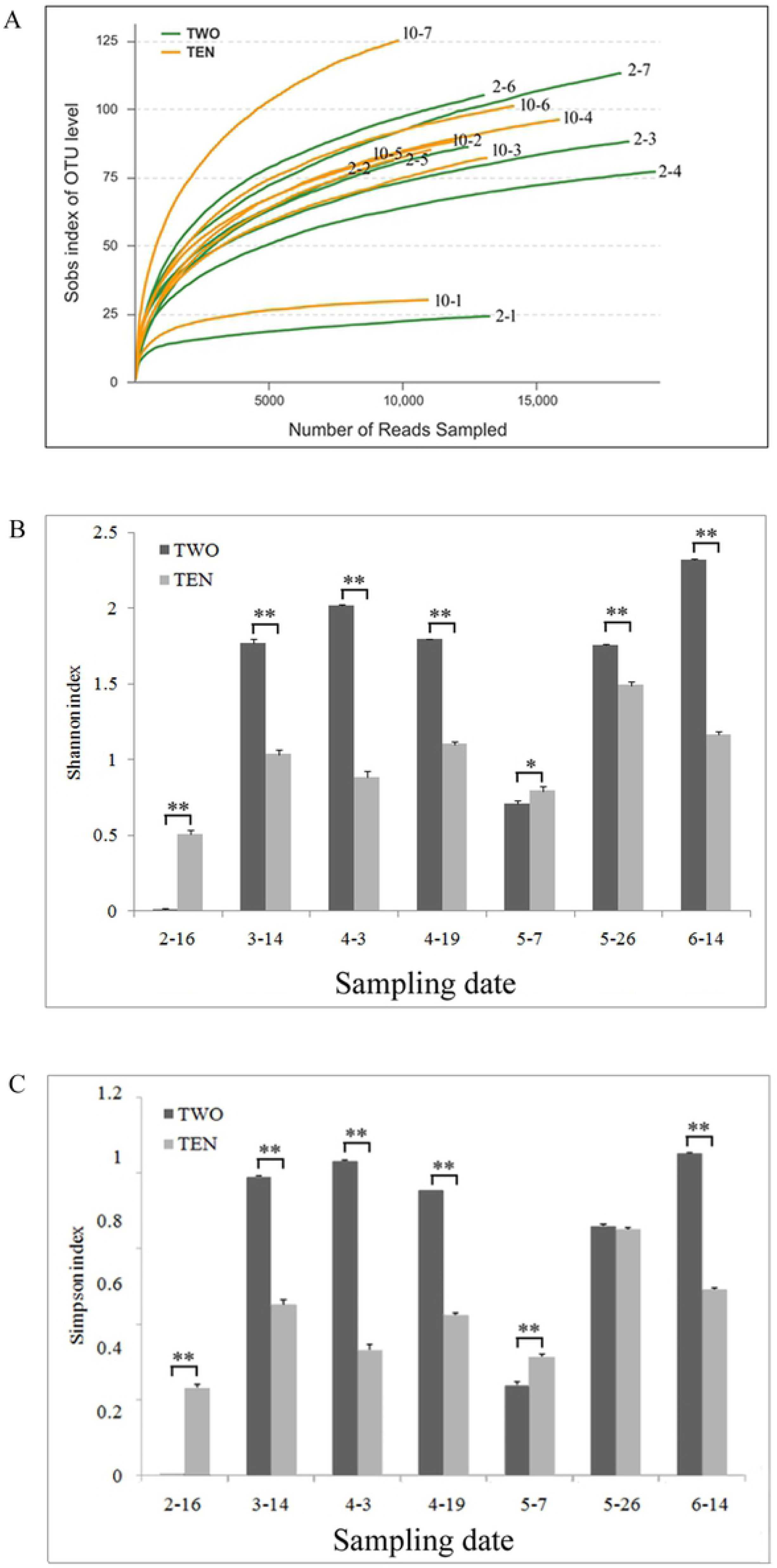
Alpha-diversity of soil nematode communities. Rarefaction curves (A). Shannon index of soil nematode communities (B). Simpson index of soil nematode communities (C). 10- was the 10-year greenhouse. 2- was the 2-year greenhouse. −1, −2, −3, −4, −5, −6, −7 were represented sampled date 2-16, 3-14, 4-3, 4-19, 5-7, 5-26 and 6-14.

Nematodes were classified into four trophic groups by feeding habits, i.e., bacterial-feeding nematodes (BF), fungal-feeding nematodes (FF), plant-parasitic nematode (PP) and omnivore-predators (OP). In the 2-year greenhouse, BF was the most predominant group from 2.16 to 6.14 with percentage from 100% to 85.7% (Fig. 2A). Likely, BF was the most dominant group from 2.16 to 5.26 in the 10-year greenhouse and showed a downward trend, whereas PP became the most group with the greatest number on 6.14. According to statistical analysis, BF in the 2-year greenhouse was markedly higher than in the 10-year greenhouse from 5.7 to 6.14. PP in the 10-year greenhouse explosively increased from 2.16 to 6.14 from 0.4% to 71.0%, but only slightly increased from 5.7 to 6.14 in the 2-year greenhouse. PP in the 10-year greenhouse was significantly higher than in the 2-year greenhouse and appeared from 5.7 to 6.14 (Fig. 2B). OP and FF were minority in the two types of greenhouses (Fig. 2C and 2D).

**Fig 2.**
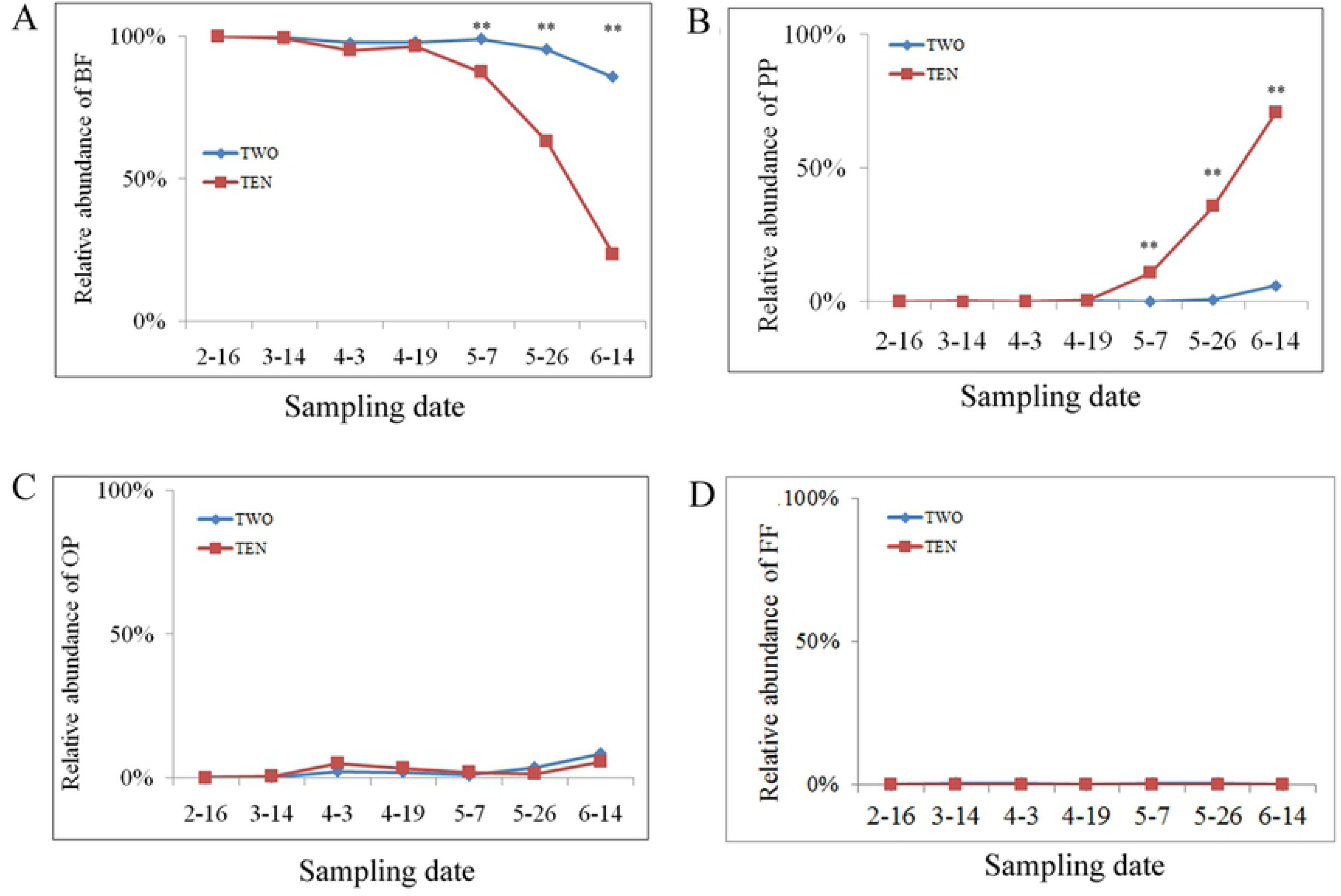
Relative abundance of trophic groups. Relative abundance of BF (A). Relative abundance of PP (B). Relative abundance of OP (C). Relative abundance of FF (D). BF, bacterial-feeding nematodes; PP, plant-parasitic nematode; OP, omnivore-predators. Blue line represented the 2-year greenhouse; Red line represented the 10-year greenhouse; *, significant at P < 0.05; **, significant at P < 0.01.

Mature index (MI) increased in the two types of greenhouses from 2-16 to 4-3, but decreased from 4-3 to 5-7. Interestingly, MI increased in the 2-year greenhouse, but continuously declined in the 10-year greenhouse from 5-7 to 6-14. The markedly differences were observed from 5-7 to 6-14 (*P*<0.01), suggesting that soil nematode communities in the two greenhouse substantially differed (Fig. 3A). Plant-parasite index (PPI) were the lowest from 2-16 to 4-3, and gradually increased in the 2-year greenhouse 4-3 to 6-14. Particularly, PPI in the 10-year greenhouse sharply increased, and significantly higher (*P*<0.01) than in the 2-year greenhouse from 5-7 to 6-14(Fig. 3B).

**Fig 3.**
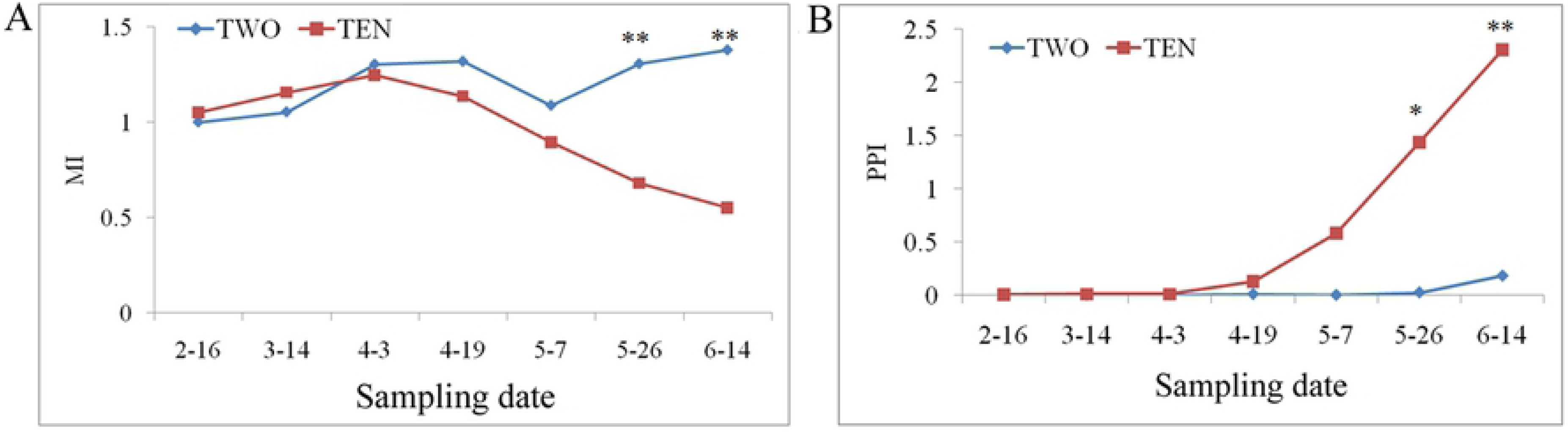
MI and PPI of soil nematode communities in the two types of greenhouses. MI (A). PPI (B). MI, maturity index. PPI, Plant parasite index. Blue line (TWO) represented the 2-year greenhouse; Red line (TEN) represented the 10-year greenhouse; *, significant at P < 0.05; **, significant at *P* < 0.01.

At order level, soil nematodes were classified into seven orders including Tylenchida, Mononchida, Plectida, Rhabditida, Triplonchida, Enoplida and Dorylaimida (Fig. 4A). Rhabditida was the predominant group (RA from 100% to 84.5%) in 7 soil samples in the 2-year greenhouse. Likely, Rhabditida was also the most dominant group (RA from 100% to 62.9%) in 6 soil samples in the 10-year greenhouse, while Tylenchida became the most abundant group (RA 71.0%). In addition, Mononchida, Plectida, Triplonchida, Enoplida and Dorylaimida were present in most soils but at relatively low abundances (RA from 0.1 to 2.1%), and unclassified soil nematode belonged to others (RA <8.4 %).

**Fig 4.**
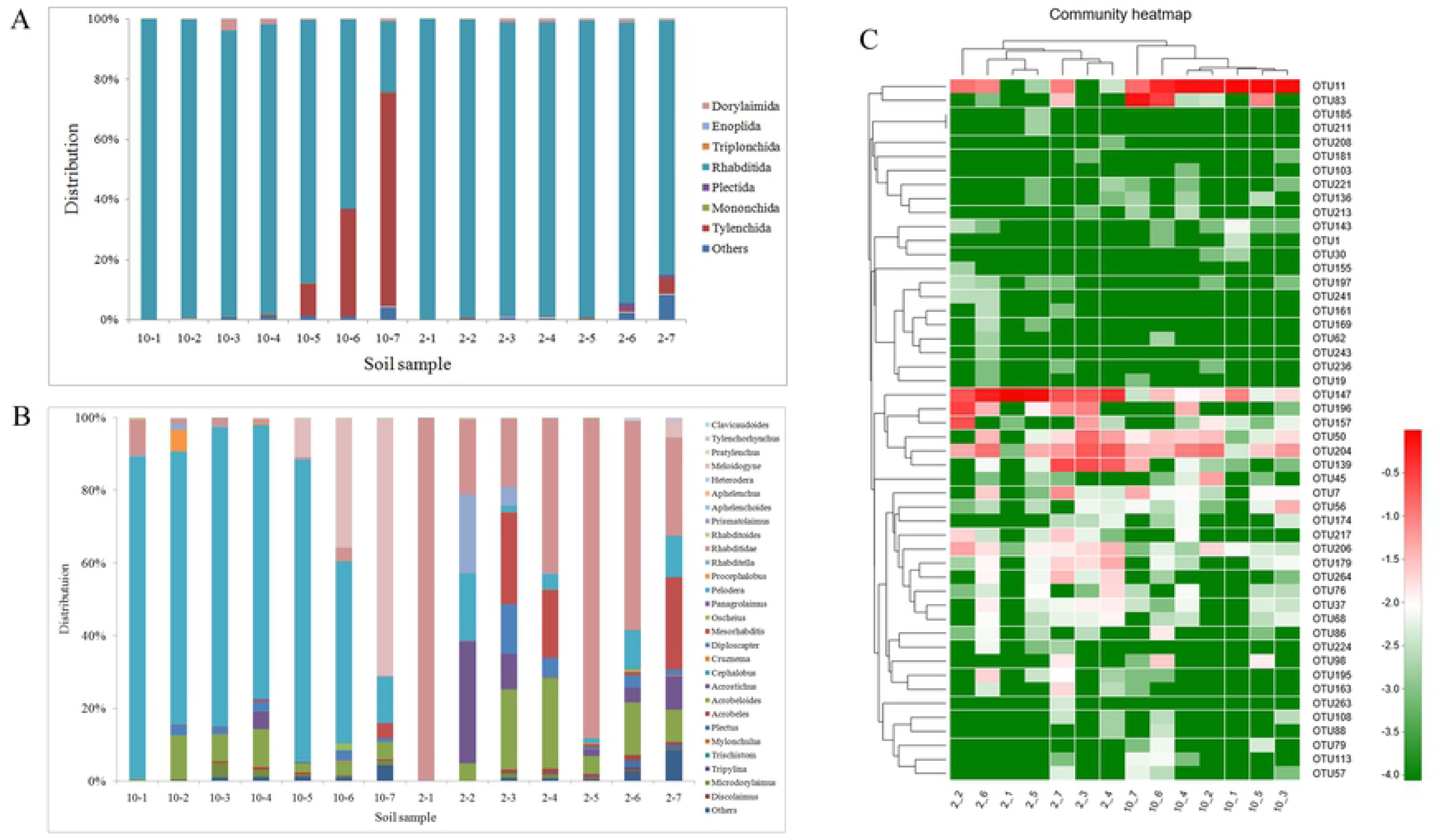
Distribution of soil nematode at order level, genus level and OUT level. Order level (A). Distribution of soil nematode at genus level (B). Distribution of soil nematode at OTU level (C). 2- was the 2-year greenhouse. −1, −2, −3, −4, −5, −6, −7 were represented sampled date 2-16, 3-14, 4-3, 4-19, 5-7, 5-26 and 6-14.

At genus level, 28 genera were observed in the two types of greenhouses (Fig. 4B). The dominant group in the 10-year greenhouse was *Pelodera* (RA from 88.6% to 50.1%) in early six samples, while *Meloidogyne* became the dominant group (RA 70.9%). However, the predominant group in the 2-year greenhouse obviously differed from the 10-year greenhouse. The most group was *Rhabditidae* (RA from 99.8% to 26.8%) in five samples (2-1, 2-4, 2-5, 2-6 and 2-7), *Acrostichus* in 2-2 sample (RA 33.6%), *Mesorhabditis* (RA 25.3%) in 2-3 sample.

A total of 50 OTUs were identified in this study. OTU11 and OTU 83 were abundant in the 10-year greenhouse, while OTU147 and OTU 139 were dominant in the 2-year greenhouse. OTU 204 and OTU 50 were both appeared with high relative abundances in the two types of greenhouses (Fig. 4C).

### Soil nematode community β-diversity

Hierarchical clustering analysis based on the Bray-Curtis distance revealed that the soil nematode communities in the two types of greenhouses differed to each other according to clustering in two big branches (Fig. 5A). UniFrac-weighted PCA analysis based on the OTU composition indicated that variations existed among these different soil nematode communities, with the first two axes explaining 66.49% and 19.53 % of the total variation (Fig. 5B). It was founded that soil nematode communities in the 2-year greenhouse were clustered to the left side in the PCA plot, while soil nematode communities in the 10-year greenhouse were clustered to the right side, indicating a clear separation among the two types of greenhouses. We also founded that soil nematode communities in one greenhouse slightly separated, indicating that soil nematode communities on various sampling time in one greenhouse had some differences, also demonstrating that soil nematode communities constantly change along with cucumber growing. The distribution of 14 soil nematode communities along the horizontal axis could indicate the changes from the 2-year greenhouse to the 10-year greenhouse. Meanwhile, the distribution of soil nematode communities along the vertical axis could reveal the changes from sampling time 2-16 to 6-14. Principal coordinate analysis (PCoA) also clearly demonstrated a similar result of the PCA with the first two axes explaining 52.84 and 16.64 % of the total variation (Fig. 5C).

**Fig 5.**
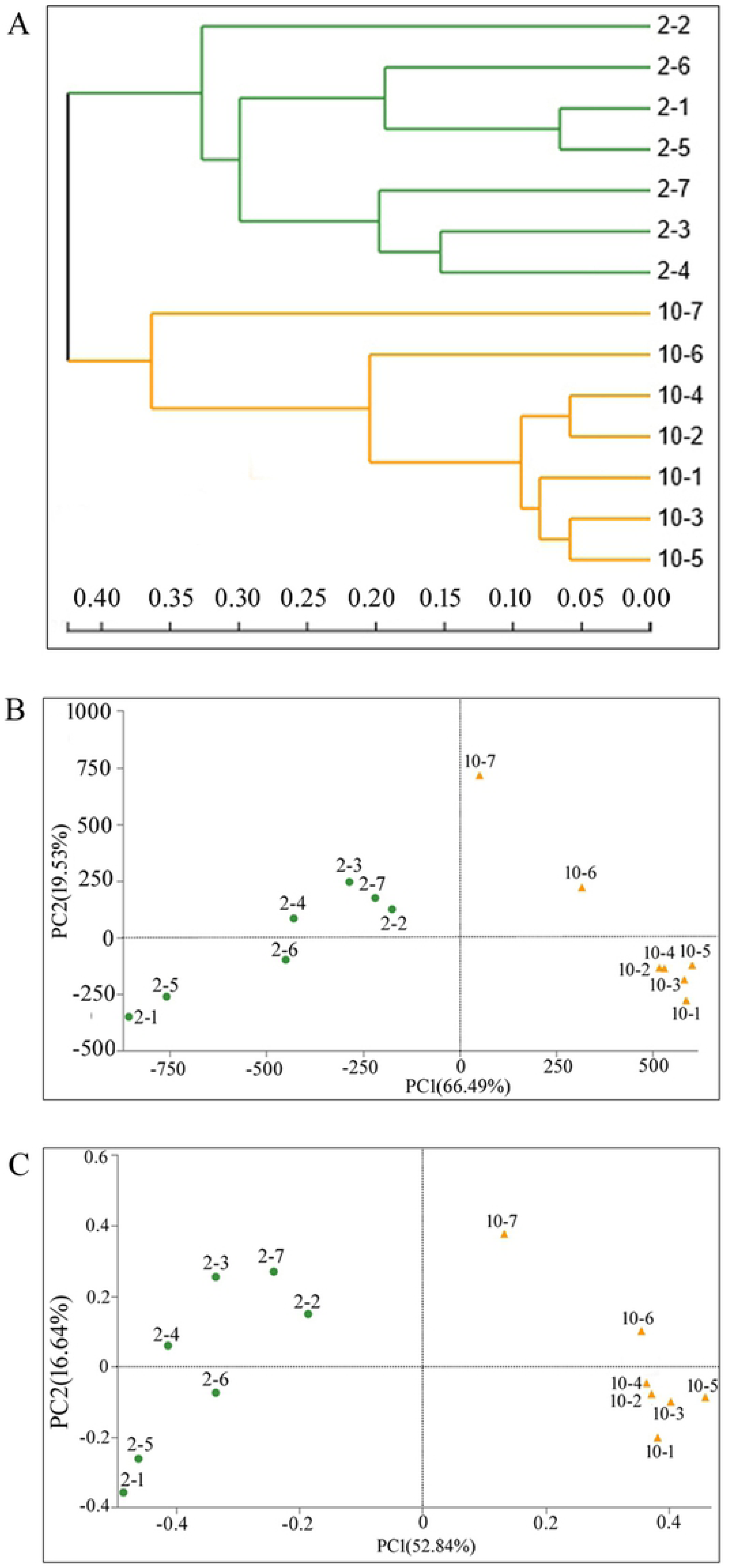
Analysis of hierarchical clustering, UniFrac-weighted PCA and PCoA. Hierarchical clustering (A). UniFrac-weighted PCA (B). PCoA plots (C). 10- was the 10-year greenhouse. 2- was the 2-year greenhouse. −1, −2, −3, −4, −5, −6, −7 were represented sampling date 2-16, 3-14, 4-3, 4-19, 5-7, 5-26 and 6-14, respectively.

In view of significant differences in later samples (10-6, 10-7and 2-6, 2-7) compared by MI, PPI, abundance of BF and PP, it was need to deeply screen which OTU caused the differences in soil nematode communities in the two types of greenhouses used by the Fisher’s exact test. It was found that OTU83 in 10-7 sample were markedly higher than in 2-7 sample with the least *P* value (Fig. 6A). OTU 11 in 10-6 sample was significantly higher than in 2-6 sample, while OTU 147 was distinctly lower than in 2-6 sample (Fig. 6B). The similar results were obtained from 10-5 and 2-5 samples. These results showed that OTU 83, OTU 11 and OTU 147 probably bring the main differences in the two types of greenhouses.

**Fig 6.**
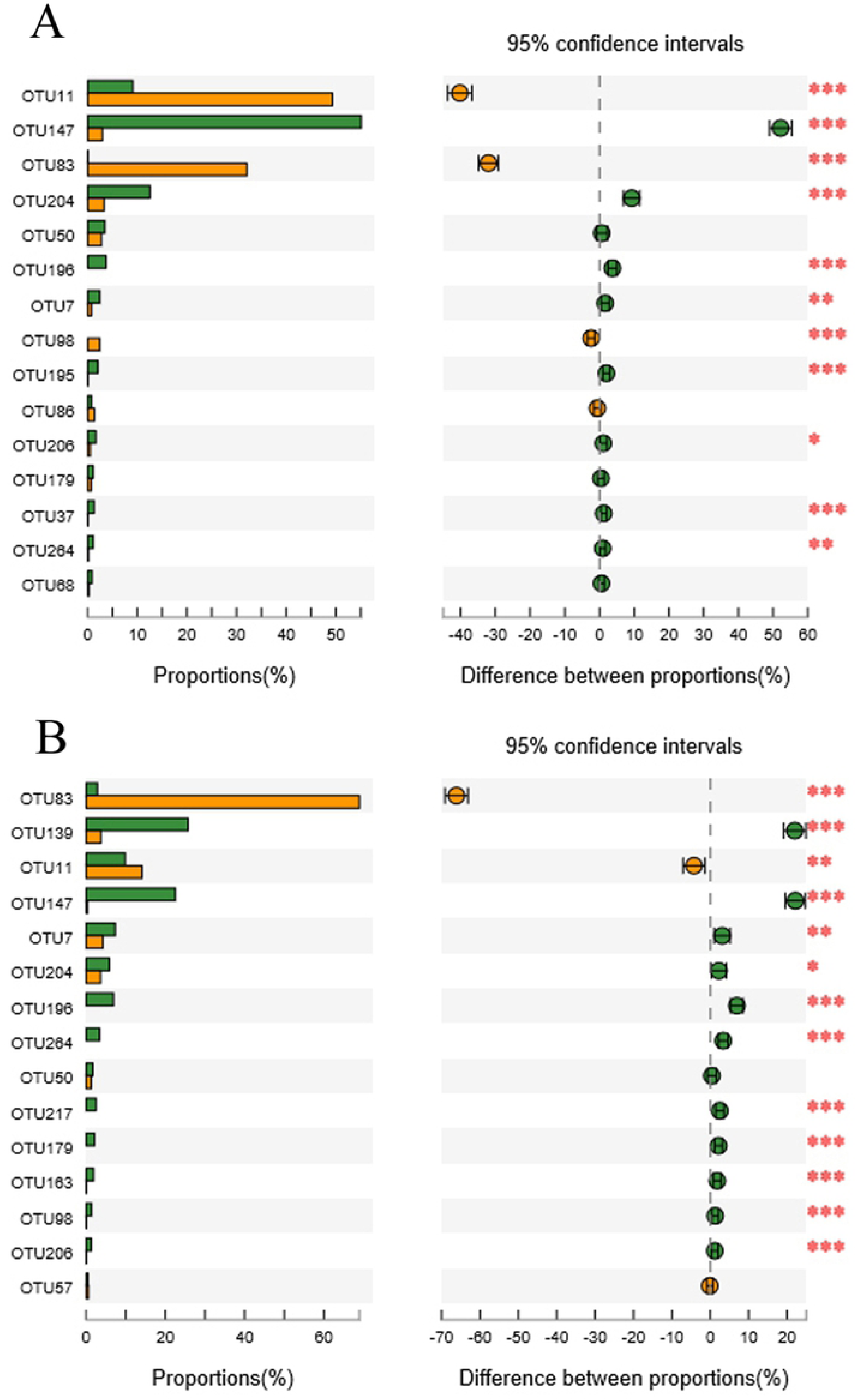
Fisher’ exact test between later samples. Test between 10-7 and 2-7(A). Test between 10-6 and 2-6 (B).Yellow represented sample in the 10-year greenhouse; Green represented sample in the 2-year greenhouse.

#### Relationships between soil nematode community, soil properties and disease index

RDA biplot (Fig. 7) was drawn to reveal the relationships between soil nematode community, soil properties and disease index of root knot nematode. It was found that high TN, SOC and EC were related to soil nematode community in the 10-year greenhouse. Two soil nematode communities (10-6 and 10-7) showed high relation to DI, suggesting that they were probably response to root knot nematode. The samples in the 2-year greenhouse were only correlated with to pH.

**Fig 7.**
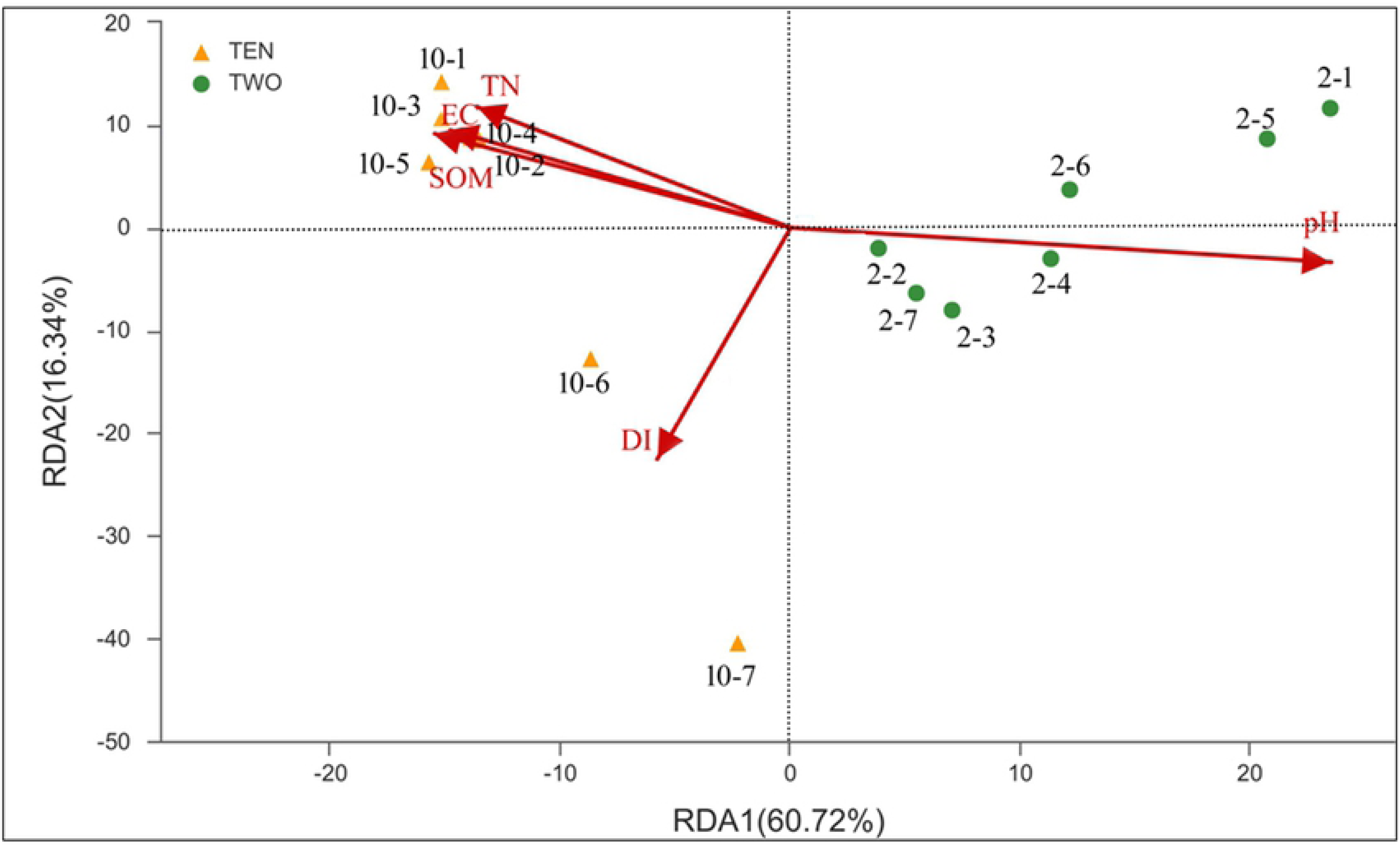
Redundancy analysis (RDA) between soil nematode communities in the two types of greenhouses and soil properties and root knot nematode disease index. SOM: soil organic matter, TN: total nitrogen, EC: electrical conductivity; DI, disease index of root knot nematode

Pearson correlation coefficients were calculated to uncover the relationship between soil nematode trophic groups, the most ten OTUs and soil properties, disease index of root knot nematode (DI) (Fig. 8). BF negatively correlated to DI (correlation coefficient −0.985, *P*<0.01), while PP positively related to DI (correlation coefficient 0.990, *P*<0.01). OTU 83 had significantly positive correlation with DI (correlation coefficient 0.987, *P*<0.01), reflecting that this OTU were responsible to the disease of root knot nematode. However, OTU11 showed positive correlation with SOM, TN and EC, negative correlation with pH, suggesting that they tend to live in high nutrients environment. OTU147 abundant in the 2-year greenhouse showed positive correlation with pH, suggesting that they probably sensitive to pH.

**Fig 8.**
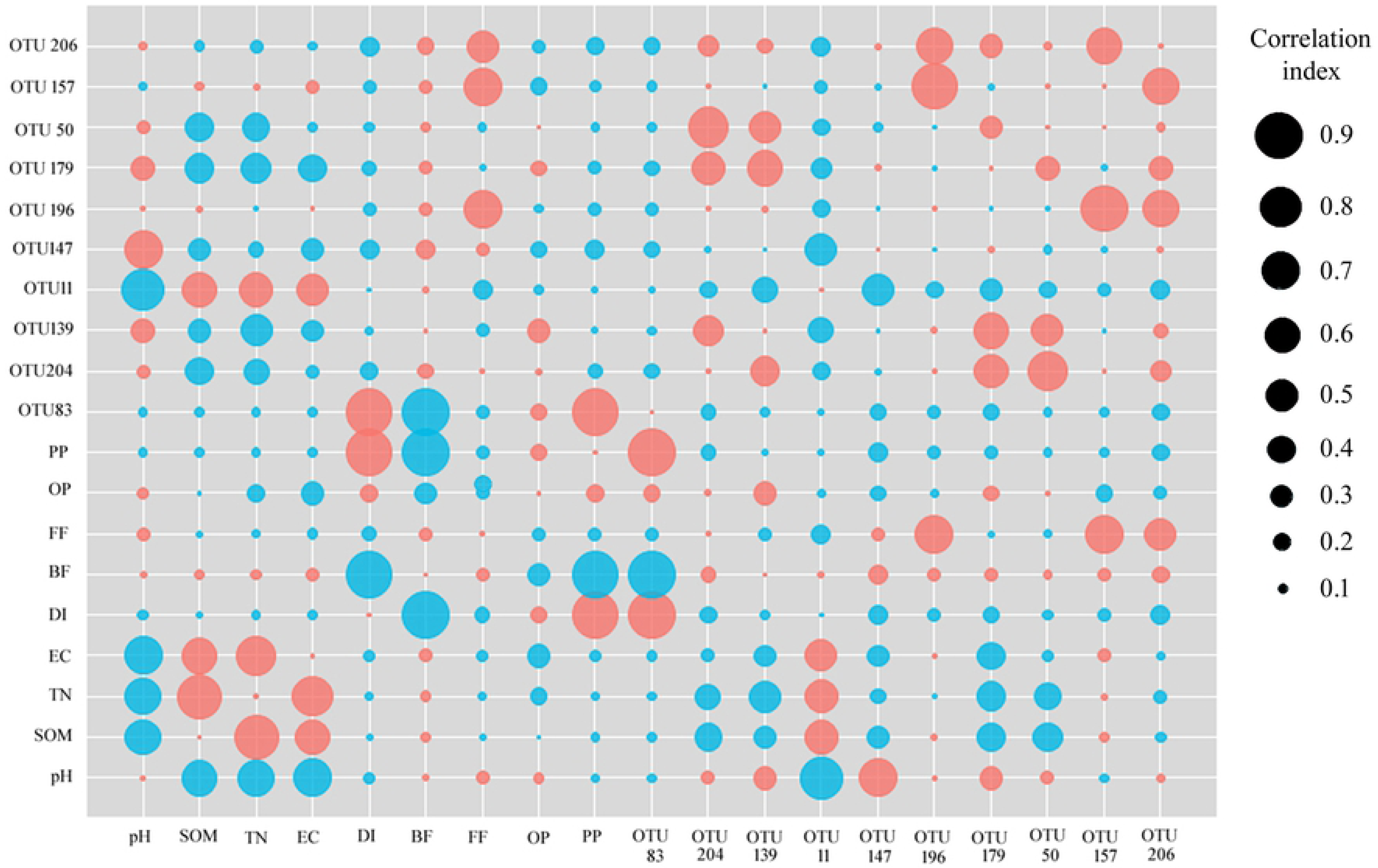
Bubble diagram of Pearson’s correlation of soil nematode trophic groups and the most ten OTUs between the soil properties and root knot nematode disease index. SOM: soil organic matter, TN: total nitrogen, EC: electrical conductivity; DI, disease index of root knot nematode. BF, bacterial-feeding nematodes; PP, plant-parasitic nematode; OP, omnivore-predators. Red bubble represented positive correlation and blue bubble represented negative correlation. The size of bubble resented correlation index.

### Population dynamics of *Meloidogyne*

We also analyzed the population dynamics of *Meloidogyne* among whole growing season to reveal accumulation of the nematode (Fig. 9). It was founded that the least abundance of *Meloidogyne* were observed from 2-16 to 4-3 with proportions from 0 to 0.2%. Subsequently, the abundance of *Meloidogyne* slowly increased from 4-3 to 5-7 with proportions from 0.3% to 10.7%, but exponentially increased from 5-7 to 6-14 with proportions from 35.5% to70.9% in the 10-year greenhouse. Hence, the population dynamics of *Meloidogyne* in the 10-year greenhouse was divided into the dormant phase, the low-level increasing phase and the exponential phase during the whole season. However, population dynamics of *Meloidogyne* in the 2-year greenhouse was only slightly increased.

**Fig 9.**
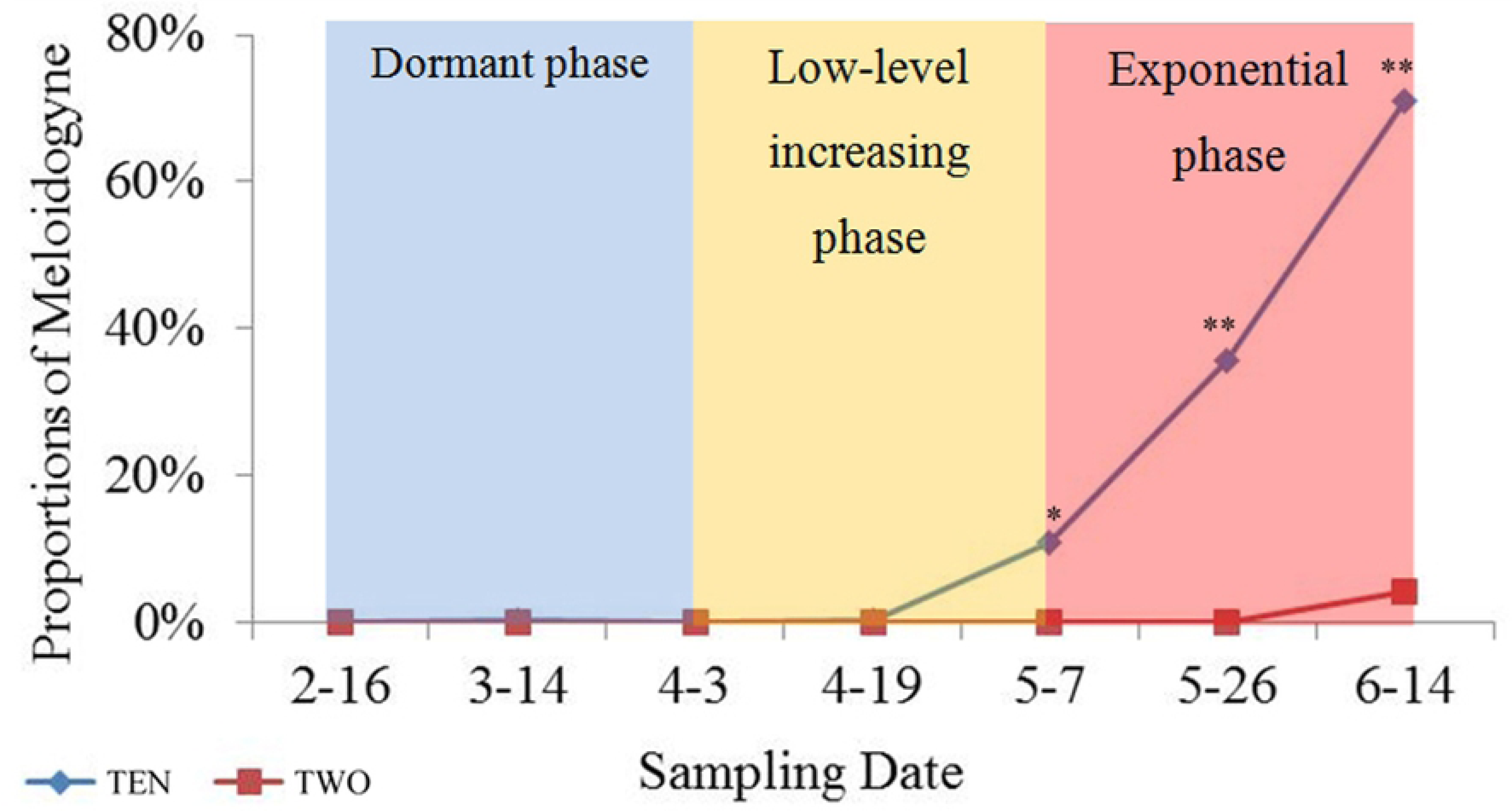
Population dynamics of *Meloidogyne* among whole growing season. Population dynamics were classified into dormant phase, Low-level increasing phase and exponential phase. Blue line (TWO) represented the 2-year greenhouse; Red line (TEN) represented the 10-year greenhouse; *, significant at *P* < 0.05; **, significant at *P* < 0.01.

## Discussion

### Differently temporal dynamic model of soil nematode communities in the two types of greenhouses

Through seven sampling times during the whole growing season of cucumber in the two types of greenhouses, we observed that soil nematode community exhibited differently dynamic change. In the 2-year greenhouse, BF was always the predominant groups and slightly decreased over the whole growing season. However, BF was dominant group on early stage and PP became the main group on late stage in the 10-year greenhouse. Many previous researches reported that BF are general the dominant nematodes in some types of soil [17,18,34]. The higher relative abundance of BF in soil was related to organic input, which led to a higher population of bacteria [35]. Most of BF have short life cycles and reproduce quickly, are quickly respond to increasing soil nutrients [36]. In this study, abundant of BF in the greenhouse indicated a relatively high nutrient content because of a plenty of organic fertilizer application. Interestingly, abundance of BF in the two greenhouse were extremely high (abundance > 90%) on early stages. The results probably derived from other two reasons: soil temperature was low when sampled early; no plants or seedling plants grow in field, thus only BF could survive. Jiang et al also found BF as the main groups in early stage [37] and can verify our results.

In our greenhouse experiment, PP became the dominant groups in the10-year greenhouse on late sampling dates, but still rare in the 2-year greenhouse (Fig. 2). The difference derived from obvious PP increasing in the 10-year greenhouse. In general, PP are not the dominant group in natural or agricultural soils, but they became the most abundant nematode in the greenhouse soil after a long period of continuous cropping [8]. In this research, PP accumulated in the 10-year greenhouse due to continuous planting. Along with temperature rising, cucumber root biomass increased and provided more feeding sties for PP, and then PP rapidly reproduced on late stage of whole growing season. Our results were similar to two reports by Korthals et al and Quist et al that PP directly depended on plant resources and closely changes with vegetation [38,39]. However, PP in the 2-year greenhouse not significantly increased because of short continuous cropping. The different temporal dynamic of soil nematode communities in the two types of greenhouses reflect that the soil nematode community had substantial discrimination between the 2-year greenhouse and the 10-year greenhouse.

### Differences in composition of soil nematodes community between the two types of greenhouses

To further analyze the differences in soil nematodes community, we compared two basic indexes (Shannon index and Simpson index) of the two types of greenhouses on 7 sampling dates. Shannon index and Simpson index of soil nematode community in the 10-year greenhouse were markedly lower than that in the 2-year greenhouse on 5 and 4 sampling times, respectively, suggesting that diversity of soil nematode community decreased with increasing planting years. Similarly, Li et al report that Shannon index of nematode fauna decrease along with increasing in strawberry continuous-cropping duration [40]. For α-diversity, the composition of soil nematode community in the two types of greenhouses significantly differed on late stage of growing season, mainly attributed to plant parasitic nematode increasing. In general, it is a common consequence of plant parasitic nematode expansion with increasing planting time [41,42]. The β-diversity analysis showed that substantial variation exists in the soil nematode communities in the two types of greenhouses by hierarchical cluster dendrograms, PCA and PCoA. The variation mainly derived from two types of greenhouses rather than various sampled time in one type of greenhouse. The soil nematode communities in total of 14 samples were classified into two groups i.e. two types of greenhouses by PCA and PCoA, also demonstrating that the main differences were derived from various greenhouses.

To screen which OTUs caused differences in the two types of greenhouse, we conducted a Fisher’ exact test between late samples and found that OTU 83, OTU 11 and OTU 147 possessed the most marked difference. OTU 83 identified as *Meloidogyne* and OTU 11 belong to *Pelodera* enriched in 10-year greenhouse. This is consistent with the fact that *M. incognita* is the dominant nematode in many greenhouses with long-term continuous cropping [18,43,44]. The result of *Pelodera* dominant in the 10-year greenhouse was similar to Mahamood’s reports that *Pelodera* was found to be having the maximum mean biomass and importance value in the whole nematode community [45]. OTU 147 merely assigned to Rhabditida was abundant in the 2-year greenhouse. Generally, Rhabditida as bacterial feeder with high percentage are usually observed from many ecological soil [46,47]. The discrimination in the dominant nematode groups in the two types of greenhouses partly reflected the differences in the soil nematode community between the 2-year greenhouse and the 10-year greenhouse.

MI and PPI can be used to distinguish differences in nematode community structure in different ages of greenhouses [18]. The MI, a measure based on the composition of the nematode community, can reflect the degree of disturbance of the soil ecosystem [30]. In our study, the 2-year greenhouse and the 10-year greenhouse bear different levels of human intervention showed by MI and PPI. MI slightly increased, subsequently declined in the 10-year greenhouse, while the PPI markedly increased. However, MI and PPI in the 2years greenhouse did not shown significantly change. Some researchers reported the inverse relationship between the MI and PPI as well as the MI decreases and PPI increases [30,48]. Our study also showed an inverse relationship between the MI and PPI. In addition, MI and PPI in the 10-year greenhouse were lower and higher than in the 2-year greenhouse from 5-7 to 6-14, reflecting that high degree of disturbance of the soil ecosystem appeared in the 10-year greenhouse.

### Differences in soil properties in the two types of greenhouses

In our study, soil properties in the 2-years and 10-year greenhouses differed significantly. High SOM, TN, EC and low pH were observed in the10-year greenhouses soil (Table 1). Likely, RDA between soil samples and soil properties also illustrated the similar results, reflecting that soil degradation such as nutrients accumulation, salinization and acidification happened in the 10-year greenhouse. This results are in line with Li et al founds that the greenhouse soil was positively correlated with SOM, TN, AK, and AP, and negatively correlated with pH [49]. In order to obtain high yield and high quality of vegetable, farmers usually applied abundant organic and N fertilizer to greenhouse filed. However, high input of nutrition were not fully utilized by soil microorganisms and plants, the extra nutrition remained in the soil, thus lead to a series of soil problems such as soil degradation, salt accumulation and pH declining in greenhouse with long planting history [13,49-51]. A general conclusion that N fertilization caused the soil pH decreased while EC increased significantly, indicating that long-term inorganic N fertilization causes soil acidification and salinization [13,52-54]. The pH decrease may be resulted from the production of acidic metabolites came from the process of urea assimilation by soil microorganisms [55] and from the process of nitrification [56]. In addition, soil properties of different sampling times in one greenhouse differed, suggesting that soil properties shifted along with cucumber growing.

Variation of soil properties could have directly or indirectly influences on soil nematode communities and lead to significantly change in structure and composition of soil nematode communities [18]. The results of Pearson correlation coefficients between soil nematode trophic groups, the most ten OTUs and the soil properties revealed that OTU11 (*Pelodera*) showed positive correlation with SOM, TN and EC, negative correlation with pH, suggesting that they tend to live in high nutrients environment. In general, high SOM cause more bacteria, thus BF would be enhanced [35].

### Dynamic population of *Meloidogyne*

The important result of our research was demonstration of dynamic population of root knot nematode. As shown in Fig. 9, the population dynamics of *Meloidogyne* was classified into the dormant phase, the low-level increasing phase and the exponential phase. It was similar to the results that no root-knot nematodes are obtained from the earlier soil samples (April), but abundant nematodes were observed later (May) [57]. During the exponential phase, the primary *Meloidogyne* infestation lay many eggs and hatched J2s, causing a secondary infestation that produced many offspring. Thus, the *Meloidogyne* population grew rapidly. In fact, *Meloidogyne* population is influenced by many abiotic and biotic factors, including organic amendments and other nematode trophic groups. Oka review the mechanism nematode suppression by organic amendments [58], but this viewpoint is contradict to our results. In our experiment greenhouse, plenty of organic amendments were fertilized to soil, while *Meloidogyne* population always increased. The possible explanation is that abundant *Meloidogyne* in soil because of long-term continuous cropping, also demonstrated that soil properties was not the main factor for root knot nematode expansion. Shi et al found that number of second stage juveniles (J2) of *Meloidogyne* sp. increases with continuous cropping years [44]. Zheng et al also reported that plant-parasitic nematodes, especially *Meloidogyne* sp., are the most abundant in 12-year-old soil [19].

## Conclusion

We obtained two different dynamic changes model of soil nematode communities in greenhouse with various continuous cropping years during the whole grow season using high-throughput sequencing. These differences derived from the discrepant composition of soil nematode in the 2-years and 10-year greenhouse. We found that *Meloidogyne* was the main factor of discrimination in pattern of soil nematode community. In addition, population of *Meloidogyne* growth was classified into the dormant phase, the low-level increasing phase and the exponential phase during the whole season. This is the first and deeply analysis of the soil nematode community in greenhouse compared with long and short duration of continuous cropping.

## Supporting information

**S1 Table. Soil properties of samples in the 2-year and the 10-year greenhouse.** SOM: soil organic matter, TN: total nitrogen, EC: electrical conductivity. 10- was the 10-year greenhouse. 2- was the 2-year greenhouse. −1, −2, −3, −4, −5, −6, −7 were represented sampling date 2-16, 3-14, 4-3, 4-19, 5-7, 5-26 and 6-14. Soil properties of sample in the two types of greenhouses were compared using ANOVA.

**S2 Table. Statistics of sequences information of samples.**

## Acknowledgments

This present study was supported by the National Natural Science Funds of China [No. 31101418], National key research & development (R&D) plan [No. 2017ZY060063], China Agriculture Research System[No.CARS-25], the Science and Technology Innovation Program of the Chinese Academy of Agricultural Sciences [No. CAAS-ASTIP-IVFCAAS].

## Author Contributions

Conceptualization: Bingyan Xie.

Data curation: Xueliang Tian.

Formal analysis: Zhenchuan Mao.

Funding acquisition: Xueliang Tian, Bingyan Xie.

Methodology: Xiaoman Zhao.

Project administration: Bingyan Xie.

Supervision: Zhenchuan Mao.

Visualization: Xueliang Tian.

Writing – original draft: Xueliang Tian.

Writing – review & editing: Bingyan Xie.

